# Therapeutic effect of CT-P59 against SARS-CoV-2 South African variant

**DOI:** 10.1101/2021.04.27.441707

**Authors:** Dong-Kyun Ryu, Rina Song, Minsoo Kim, Young-Il Kim, Cheolmin Kim, Jong-In Kim, Ki-Sung Kwon, Aloys SL Tijsma, Patricia M Nuijten, Carel A van Baalen, Tandile Hermanus, Prudence Kgagudi, Thandeka Gwete-Moyo, Penny L Moore, Young Ki Choi, Soo-Young Lee

## Abstract

The global circulation of newly emerging variants of SARS-CoV-2 is a new threat to public health due to their increased transmissibility and immune evasion. Moreover, currently available vaccines and therapeutic antibodies were shown to be less effective against new variants, in particular, the South African (SA) variant, termed 501Y.V2 or B.1.351. To assess the efficacy of the CT-P59 monoclonal antibody against the SA variant, we sought to perform as *in vitro* binding and neutralization assays, and *in vivo* animal studies. CT-P59 neutralized B.1.1.7 variant to a similar extent as to wild type virus. CT-P59 showed reduced binding affinity against a RBD (receptor binding domain) triple mutant containing mutations defining B.1.351 (K417N/E484K/N501Y) also showed reduced potency against the SA variant in live virus and pseudovirus neutralization assay systems. However, *in vivo* ferret challenge studies demonstrated that a therapeutic dosage of CT-P59 was able to decrease B.1.351 viral load in the upper and lower respiratory tracts, comparable to that observed for the wild type virus. Overall, although CT-P59 showed reduced *in vitro* neutralizing activity against the SA variant, sufficient antiviral effect in B.1.351-infected animals was confirmed with a clinical dosage of CT-P59, suggesting that CT-P59 has therapeutic potential for COVID-19 patients infected with SA variant.

**Highlights:**

- CT-P59 significantly inhibit B.1.1.7 variant to a similar extent as to wild type virus
- CT-P59 showed reduced potency against the B.1.351 variant in *in vitro* studies
- Therapeutic dosage of CT-P59 showed *in vivo* neutralizing potency against B.1.351 in ferret challenge study.

## 1. Introduction

With the emergence of new variants of SARS-CoV-2, the morbidity and mortality of COVID-19 has been rising again in many countries [1-6]. For instance, the B.1.1.7 (501Y.V1) variant first described in the UK, and containing a N501Y mutation in the spike protein has spread to more than 100 countries worldwide. Currently, B.1.1.7 has already become the most prevalent virus in European countries such as Germany, France, Italy, and Spain as well as the UK. The second variant of concern (VOC), B.1.351 (501Y.V2) first discovered in South Africa variant, has 9 mutations (L18F, D80A, D215G, Δ242-244, K417N, E484K, N501Y, D614G, and A701V) [7]. This variant dominates the South African epidemic, and is now circulating to neighboring nations in Africa, and local transmission has been observed in European nations and US. Lastly, P.1 (501Y.V3) [8] detected in Brazil variant and which shares several mutations with B.1351, has since been detected in several other countries in South America. Currently, while several vaccines and therapeutics against COVID-19 have been developed, several studies indicate compromised activity against new variants, compared to original SARS-CoV-2 virus [9-13]. Neutralization assays with serum from vaccinees demonstrated that viral vectored vaccines and two mRNA-based vaccines maintained activity against the UK variant, but showed approximately 10-fold reduction against the SA variant, compared to wild type virus. Monoclonal therapeutic antibodies forming part of the REGN-10933/REGN-10987 cocktail showed neutralizing activity against UK and SA variant in *in vitro* neutralization assay. In contrast, LY-CoV555 showed preserved activity against the UK variant but activity against the SA variant was completely abolished since LY-CoV555 cannot bind to triple mutant protein (K417N/E484K/N501Y) of RBD in SARS-CoV-2 SA variant[14, 15].

We have developed a COVID-19 therapeutic antibody, CT-P59 which binds to RBD of SARS-CoV-2, interfering with binding to ACE2 (Angiotensin-Converting Enzyme 2), the cellular receptor for viral entry. Also, we have demonstrated that CT-P59 has high potency against the original variant in *in vitro* and *in vivo* studies [16]. Here, we evaluated the efficacy of CT-P59 against SA variant via both *in vitro* binding and neutralization assay with live and pseudoviruses and *in vivo* challenge experiment in ferrets.

## 2. Materials and methods

### 2.1. Cells and viruses

VeroE6 cells (ATCC, CRL-1586) were cultured in Dulbecco’s modified Eagle’s medium (DMEM) supplemented with 10% (v/v) fetal bovine serum (FBS) and 2 mM penicillin-streptomycin (100 U/mL). BetaCoV/Munich/BavPat1/2020 (European Virus Archive Global #026V-03883) was kindly provided by Dr. C. Drosten. UK variant (USA/CA_CDC_5574/2020, NR-54011) and SA variant (Isolate hCoV-19/South African/KRISP-K005325/2020, NR-54009) were obtained through BEI Resources. For *in vivo* study, NMC-nCoV02 (S clade) and hCoV-19/Korea/KDCA55905/202 (B.1.351) were provided by National Culture Collection for Pathogens.

### 2.2. Biolayer interferometry (BLI)

Binding affinity of CT-P59 to wild type and mutant SARS-CoV-2 RBDs were measured by biolayer interferometry (BLI) using the Octet QK^e^ system (ForteBio) as described previously [17]. All of the mutant SARS-CoV-2 RBDs (K417N, E484K, N501Y and triple mutant) were purchased from Sino Biological.

### 2.3. ELISA

ELISA was performed as described previously [17]. In brief, Recombinant RBD or triple mutant proteins were coated onto 96-well, high-binding plates. Following blocking, serially diluted CT-P59 were incubated, followed by an anti-human horseradish peroxidase-conjugated antibody. The signal was developed with TMB substrate (Thermo Fisher Scientific) and absorbance at 450 nm was measured.

### 2.4. Microneutralization (ViroSpot) assay

To assess the susceptibility of SARS-CoV-2 variants, a microneutralization assay was performed which was based on the ViroSpot technology described earlier [18]. Briefly, approximately 100 infectious units of virus was mixed with ten 3-fold serial dilutions CT-P59. The virus/CT-P59 mixture was incubated for 1 h prior to addition to the Vero E6 cells. After a subsequent 1 h incubation, the inoculum was removed, and 1% carboxymethylcellulose (CMC) overlay was added. The plates were incubated for a duration of 16-24 hours at 37°C and 5% CO_2_. The cells were formalin-fixed and ethanol permeabilized followed by incubation with a murine monoclonal antibody which targets the viral nucleocapsid protein (Sino Biological), followed by a secondary anti-mouse IgG peroxidase conjugate (Thermo Scientific) and TrueBlue (KPL) substrate. This formed a blue precipitate on nucleocapsid-positive cells. Images of all wells were acquired by a CTL Immunospot analyzer, equipped with Biospot^®^ software to quantitate the nucleocapsid-positive cells (= virus signal). The IC50 was calculated according to the method described earlier [19].

### 2.5. Pseudovirus assay

Pseudovirus assay was performed as described previously [17]. In short, pseudovirus and serially diluted CT-P59 were incubated and the cells were added. After incubation, luminescence was measured using PerkinElmer Life Sciences Model Victor X luminometer. Neutralization was measured as described by a reduction in luciferase gene expression after single-round infection of 293T/ACE2.MF cells with spike-pseudotyped viruses.

### 2.6. Animal experiments

The animals were housed in BSL3 facility within Chungbuk National University (Cheongju, Korea) with 12 h light/dark cycle with supplied to water and food. All animal cares were performed strictly following the animal care guideline and experiment protocols approved by Institutional Animal Care and Use Committee (IACUC) in Chungbuk National University (CBNUA-1473-20-01)

### 2.7. Ferret study

Groups of 14- to 16-month-old female ferrets (Mustela putorius furo, n=6/group) were sourced from ID.BIO, Corp., Korea. All ferrets, which are seronegative for SARS-CoV-1 and SARS-CoV-2, were inoculated intranasally and intratracheally with 10^5.5^ TCID_50_ of wild type and B.1.351, respectively, while under anesthesia. Two doses of CT-P59, 80 mg/kg and 160 mg/kg, were administered intravenously 24 h after virus inoculation in treatment groups. Animals in the control group were given vehicle control. Viral load was measured from nasal wash and lung as the representative for upper and lower respiratory tract. Body weight was daily measured and clinical signs were observed throughout the study. Cough, rhinorrhea and movement were scored and analyzed for each animal in blind manner.

### 2.8. Virus titration and quantitation

The viral loads in nasal washes and lungs were determined via qRT-PCR and TCID_50_ as previously described [16]. Briefly, Viral RNA was extracted and cDNAs were generated. Viral copy numbers were determined by real-time qRT–PCR with SARS-CoV-2-specific primer set. Real-time PCR reactions were performed using a SYBR Green Supermix (Bio-Rad) and a CFX96 Touch Real-Time PCR Detection System (Bio-Rad). Viral titers for samples from the upper and lower airway were quantitated via TCID_50_ which is determined according to the method of Reed and Muench [20]. Virus titer and viral RNA from animals were analyzed using GraphPad Prism v 9.1.0.

## 3. Results

### 3.1. CT-P59 showed reduced potency against SA variant in *in vitro* assays

To assess the efficacy of CT-P59 against SA variant, we determined the binding affinity of CT-P59 against individual and triple mutant RBDs (K417N, E484K, N501Y, and K417N/E484K/N501Y), which define the SA variant, by using BLI. The equilibrium dissociation constant (K_D_) of CT-P59 against K417N was equivalent to that against wild type. K_D_ values of CT-P59 were reduced by approximately 2-, 2-, and 10-fold in E484K, N501Y, and triple mutant (K417N/E484K/N501Y), respectively (Table 1). In addition, ELISA showed that CT-P59 had reduced binding affinity against a triple mutant RBD, compared to wild type (Fig. 1A). To evaluate whether CT-P59 can neutralize B.1.351, we performed two kinds of cell-based assays using either live virus or pseudotyped virus. The live virus neutralization assay showed approximately 20-fold reduced susceptibility against SA variant, compared to wild type and UK variant (Fig. 1B and Table 2). Similarly, CT-P59 inhibited SARS-CoV-2 D614G pseudovirus with IC_50_ value of 10 ng/mL, but showed 33-fold reduced neutralization of B.1.351 pseudotyped viruses, with an IC_50_ value of 330 ng/mL (Fig. 1C and Table 3). Taken together, while CT-P59 showed reduced neutralization of B.1.351, remaining binding and neutralizing activities against B.1.351 provided a strong rationale to study the antiviral efficacy of CT-P59 in a relevant animal model whether it is affected by the reduced potency observed *in vitro*.

**Table 1.**
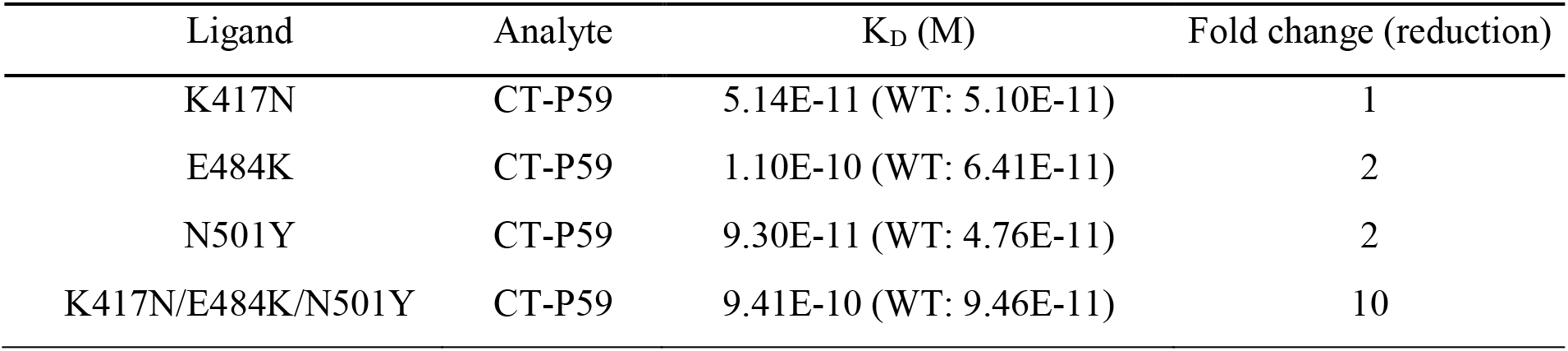
Binding affinity between CT-P59 and SARS-CoV-2 RBD mutants determined by BLI.

**Table 2.** Neutralization effect of CT-P59 against live SARS-CoV-2 variants and wild type virus.

| Viruses | Geometric mean IC<sub>50</sub> (ng/mL) | Fold change (reduction) |
| --- | --- | --- |
| BavPat1/2020 (D614G) | 2.0 | 1 |
| B.1.1.7 (UK variant) | 2.8 | 1.4 |
| B.1.351 (SA variant) | 39.5 | 19.8 |

**Table 3.** Neutralization effect of CT-P59 against SA variants and wild type pseudoviruses.

| Viruses | IC<sub>50</sub> (ng/mL) | Fold change (reduction) |
| --- | --- | --- |
| D614G | 10 | 1 |
| B.1.351 (SA variant) | 330 | 33 |

**Figure 1.**
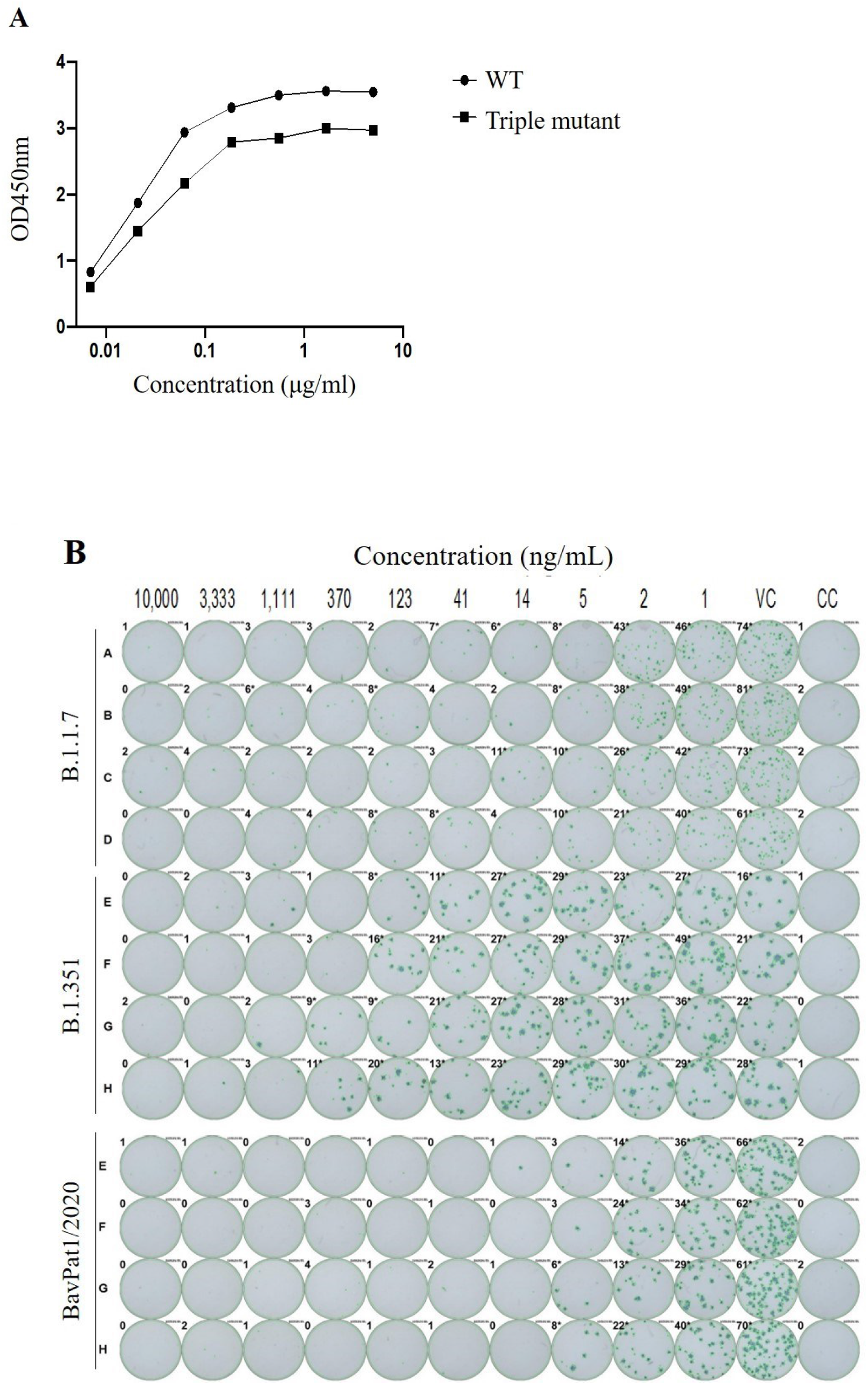
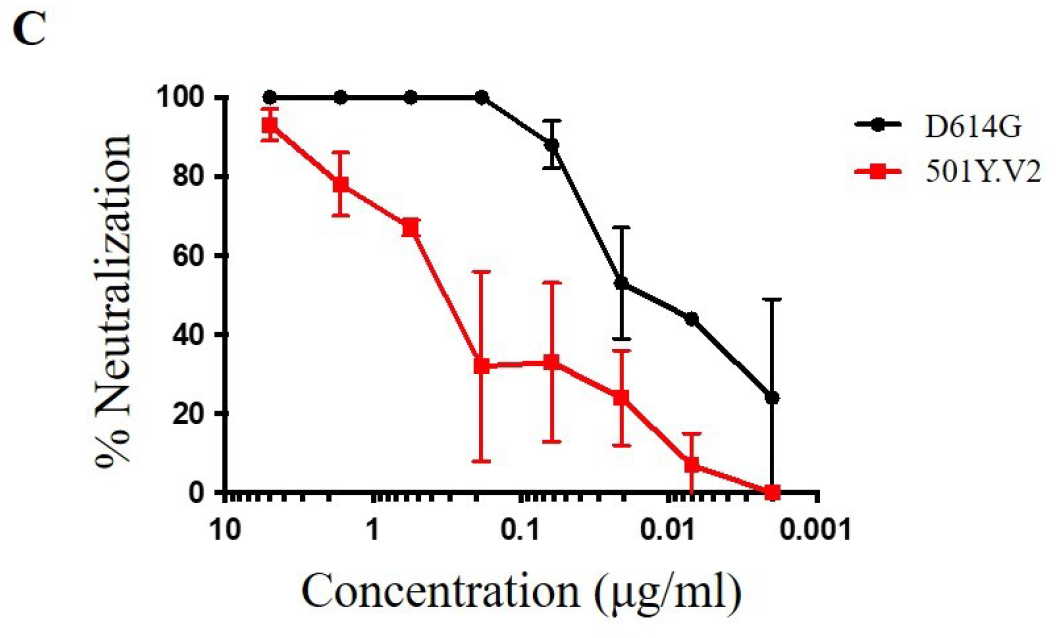
*In vitro* binding and neutralization of CT-P59 against B.1.351. Wild type RBD (circle) or SA triple RBD mutant protein (rectangle) was coated and incubated with CT-P59, followed by optical density (OD) measurement. Binding affinity of CT-P59 was measured by ELISA (A). CT-P59 was premixed with SARS-CoV-2 variant and wild type viruses (B.1.1.7, B.1.351, and BavPat1/2020). The inocula were added into VeroE6 cells, incubated and detected by anti-nucleocapsid antibody and staining (B). VC and CC represent as virus control and cell control, respectively. In addition, serially diluted CT-P59 mixed with D614G (black circle) or 501Y.V2 (red rectangle) variant pseudovirus. %Neutralization was calculated by measuring luciferase activities.

### 3.2. *In vivo* efficacy in animal models

To demonstrate the *in vivo* antiviral efficacy of CT-P59 against B.1.351, we conducted virus challenge studies in the ferret animal model. Dose of 160 mg/kg for ferret was selected as clinically relevant dose by conversion from the clinical dose of 40 mg/kg based on the identical exposure to CT-P59 in ferrets. Then, 80 mg/kg was additionally selected as a half dose. Since the initial exposure to therapeutic antibody during the early period of treatment is important factor to prevent from the worsening of infectious disease [21], the dose conversion between human and ferret was based on the exposure for first three days (AUC_0-72hr_) (data not shown).

The animals challenged with B.1.351 showed significantly reduced viral RNA in nasal washes and lung following CT-P59 treatment (Fig. 2 A and B). In nasal washes, viral RNA at both 80 mg/kg and 160 mg/kg of doses were decreased by 2.97 log_10_/mL compared to the control one day after CT-P59 administration (2 dpi). In lungs, even though viral RNA of all animals including control animals at 4 dpi had declined to BLQ (below the limit of quantitation), there was a significant decrease in both treatment groups at 2 dpi (1.30 log and 1.57 log reduction for 80 mg/kg and 160 mg/kg, respectively) compared to the control group.

**Figure 2.**
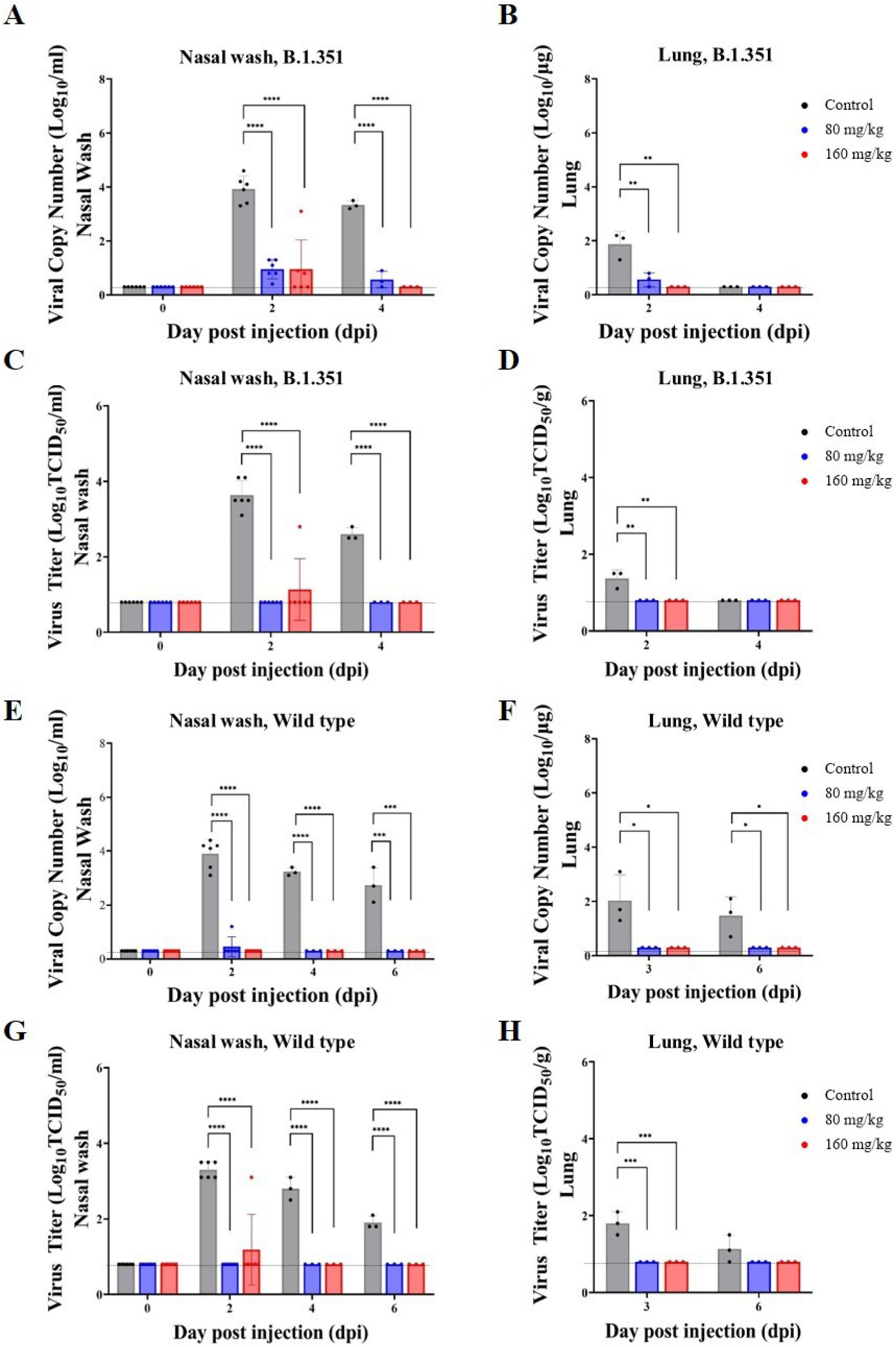
*In vivo* efficacy of CT-P59 against B.1.351 and wild type virus in animal model. Female ferrets (n=6/group) were challenged with 10^5.5^ TCID_50_/ml of B.1.351 or wild type virus, and CT-P59 was administered intravenously after 24 hours of virus inoculation. Both viral RNA (A, B, E, F) and viral titer (C, D, G, H) were measured from nasal wash and lung by using qPCR and TCID_50_, respectively. The graphs are shown as means ± SD from six or three animals at each interval, and dashed lines indicate below the limit of quantification (BLQ) as following; 0.3 log_10_ copies Log_10_/mL (Nasal wash) and 0.3 log copies/mL (Lung) for qPCR, 0.8 log_10_ TCID_50_/mL (Nasal wash) and 0.8 log_10_ TCID_50_/g (Lung) for TCID_50_ assay. Asterisks indicate statistical significance between the control and each group as determined by one-way ANOVA and post-hoc Dunnett’s test. (* indicates *P* < 0.05, ** indicates *P* < 0.01, *** indicates *P* < 0.001, and **** indicates *P* < 0.0001).

The infectious virus titer (TCID_50_) in nasal washes and lungs was also significantly decreased in animals treated with CT-P59 when compared to controls, and the infectious virus of B.1.351 was not detected in nasal washes from 2 dpi, with the exception of one animal in the 160 mg/kg treatment group at 2 dpi (Fig. 2 C and D). In lungs, virus was not detected from 2 dpi in both treatment groups, and the virus titer of control group was also not detected at 4 dpi. As expected, the animals challenged with wild type virus also showed significant reduction in viral RNA following CT-P59 treatment in nasal washes, and in lung (Fig.2 E and F). The infectious virus titer (TCID_50_) in nasal wash and lung was also significantly decreased in animals treated with CT-P59 when compared to controls (Fig. 2 G and H).

Ferrets challenged with either B.1.351 or wild type showed a decrease in body weight by about 5% in the control group, while there was no weight loss in the CT-P59 administered groups (Supplementary Fig. 1). There were minimal to mild clinical symptoms after CT-P59 administration until 3dpi; however, no clinical signs were observed from 4 dpi, except one animal challenged with SA variant (Supplementary Table 3 and 4).

## 4. Discussion

Here, we have demonstrated the therapeutic potential of neutralizing antibody CT-P59 against the B.1.351 variant of SARS-CoV-2 in *in vitro* and *in vivo* studies. Unlike LY-CoV555, CT-P59 was able to bind to RBD of B.1.351and neutralized B.1.351, albeit with reduced K_D_ and IC_50_ values. This is a suggestion that CT-P59 showed high barrier to resistant viruses since CT-P59 has relatively conserved residues for RBD binding although K417 and E484 are part of the epitope of CT-P59 [16].

Also, we demonstrated the *in vivo* neutralizing potency of CT-P59 in clinically relevant doses against B.1.351 and wild type virus in the ferret model. It was reported that SARS-CoV-2 preferentially replicates in the upper respiratory tract of ferrets, thus the relatively lower virus titer in the lower respiratory tract is likely in line with mild symptoms with SARS-CoV-2, which means it could represent of patients with asymptomatic to mild symptoms [22, 23]. Such findings were consistent with the previous literature [16], thereby, the current study employed ferrets for the disease model, taking the intended indication of CT-P59 for treating patients with the mild symptoms into account. As expected, ferrets showed the lower virus titer in the lower airway than the upper airway, for the wild type virus. Importantly, viral load of B.1.351 was also lower in the lower airway than the upper airway, and it decreased relatively faster than wild type virus among control groups.

The tendency of viral loads was mostly consistent in viral RNA copy and infectious virus quantitation in the respiratory tract. The viral loads expectedly decreased from one day after CT-P59 administration in ferrets with wild type virus, as previously described in [16]. CT-P59 demonstrated its *in vivo* neutralization potency against B.1.351, and trend in viral load were similar to the wild type virus. More animals showed the level of viral RNA above BLQ from the nasal washes at 2 dpi and 4 dpi from B.1.351 infected animals particularly with 80 mg/kg of dose, this slight delay in viral RNA clearance of B.1.351 might result from the reduced neutralizing activity of CT-P59 during the early phase following CT-P59 administration [24]. Nonetheless, viable virus were not detected across the entire respiratory tract from 2 dpi in the animals given 80 mg/kg, by using TCID_50_ assay. These data suggest that CT-P59 is unlikely to compromise the therapeutic effect of CT-P59 against B.1.351 in the respiratory tract in *in vivo*. This preserved capacity to neutralize B.1.351 is in contrast to complete loss of neutralization potency previously reported for commercialized therapeutic antibody [14], LY-CoV555. Moreover, *in vitro* neutralization potency of antibodies did not directly correlate with *in vivo* neutralizing effect, in terms of *in vivo* environment and Fc functionality [25-27]. Overall, considering that ferrets were administered with clinically relevant doses of CT-P59, it is expected CT-P59 would maintain its efficacy with patients infected with SA variant.

## Conclusion

The *in vitro* studies demonstrated that CT-P59 showed reduced activity against B.1.351, but not B.1.1.7 variant. Nonetheless, *in vivo* challenge studies showed reduction in viral load in B.1.351-infected ferrets by sufficient therapeutic dosage, despite the reduced potency of CT-P59 against SA variant.

## Declaration of competing interest

The authors declare that they have no known competing financial interests or personal relationships that could have appeared to influence the work reported in this paper

## Acknowledgements

We acknowledge funding from the South African Medical Research Council Strategic Health Innovation Program. P.L.M. is supported by the South African Research Chairs Initiative of the Department of Science and Innovation and the NRF (Grant No 9834)

## Supplementary Material

**Supplementary Table 1.** Wild type RBD.

| **Dilution** | **CT-P59** | **Class II mAb 1** | **Class II mAb 2** | **Class II mAb 3** | **CR3022 (positive control)** | **Palivizumab (negative control)** |
| --- | --- | --- | --- | --- | --- | --- |
| 5.000 | 3.55 | 3.33 | 3.40 | 2.58 | 3.57 | 0.00 |
| 1.667 | 3.56 | 3.49 | 3.52 | 2.28 | 3.53 | 0.00 |
| 0.556 | 3.50 | 3.41 | 3.46 | 1.28 | 3.34 | 0.00 |
| 0.185 | 3.31 | 3.26 | 3.32 | 0.52 | 3.07 | 0.00 |
| 0.062 | 2.94 | 2.46 | 2.59 | 0.07 | 2.49 | 0.00 |
| 0.021 | 1.87 | 1.55 | 1.80 | 0.03 | 1.20 | 0.00 |
| 0.007 | 0.83 | 0.67 | 0.87 | 0.01 | 0.43 | 0.00 |

**Supplementary Table 2.** Triple mutant RBD.

| **Dilution** | **CT-P59** | **Class II mAb 1** | **Class II mAb 2** | **Class II mAb 3** | **CR3022 (positive control)** | **Palivizumab (negative control)** |
| --- | --- | --- | --- | --- | --- | --- |
| 5.000 | 2.97 | 0.00 | 0.00 | 0.00 | 3.32 | 0.00 |
| 1.667 | 3.00 | 0.00 | 0.00 | 0.00 | 3.36 | 0.00 |
| 0.556 | 2.85 | 0.00 | 0.00 | 0.00 | 3.24 | 0.00 |
| 0.185 | 2.79 | 0.00 | 0.00 | 0.00 | 3.02 | 0.00 |
| 0.062 | 2.17 | 0.02 | 0.00 | 0.02 | 2.42 | 0.00 |
| 0.021 | 1.45 | 0.03 | 0.00 | 0.00 | 1.25 | 0.00 |
| 0.007 | 0.61 | 0.00 | 0.00 | 0.00 | 0.49 | 0.00 |

**Supplementary Table 3.** Clinical scores of ferrets infected with B.1.351 SARS-CoV-2.

| Group |  | 0dpi(n=6) | 1dpi(n=6) | 2dpi(n=6) | 3dpi(n=3) | 4dpi(n=3) |
| --- | --- | --- | --- | --- | --- | --- |
| **B.1.351**<br>**Infection** | Cough | 0.00 | 0.00 | 0.50 | 0.33 | 0.33 |
|  | Rhinorrhea | 0.00 | 0.00 | 0.00 | 0.33 | 0.00 |
|  | Movement, activity | 0.00 | 0.00 | 1.67 | 1.00 | 1.00 |
|  | Total | 0.00 | 0.00 | 2.17 | 1.67 | 1.33 |
| **B.1.351**<br>**80 mg/kg** | Cough | 0.00 | 0.00 | 0.67 | 0.33 | 0.00 |
|  | Rhinorrhea | 0.00 | 0.00 | 0.00 | 0.00 | 0.00 |
|  | Movement, activity | 0.00 | 0.00 | 1.00 | 0.67 | 0.00 |
|  | Total | 0.00 | 0.00 | 1.67 | 1.00 | 0.00 |
| **B.1.351**<br>**160 mg/kg** | Cough | 0.00 | 0.00 | 0.00 | 0.00 | 0.00 |
|  | Rhinorrhea | 0.00 | 0.00 | 0.00 | 0.33 | 0.00 |
|  | Movement, activity | 0.00 | 0.00 | 0.83 | 0.67 | 0.33 |
|  | Total | 0.00 | 0.00 | 0.83 | 1.00 | 0.33 |

**Observational clinical symptoms:** Cough, rhinorrhea, movement, and activity.
Score: 0; normal, 1: occasional, mild reduced activity, 2: frequent, reduced activity.
\* Scores were measured by observation of clinical symptoms for at least 20 minutes in each group of ferrets based on the following criteria
**Cough:** 0; no evidence of cough, 1; occasional cough, 2; frequent cough (score 2).
**Rhinorrhea:** 0; no nasal rattling or sneezing, 1; moderate nasal discharge on external nares, 2; severe nasal discharge on external nares.
**Movement:** activity: 0; normal movement and activity, 1; mild reduced movement and activity, 2; evidence of reduced movement and activity.

**Supplementary Table 4.**
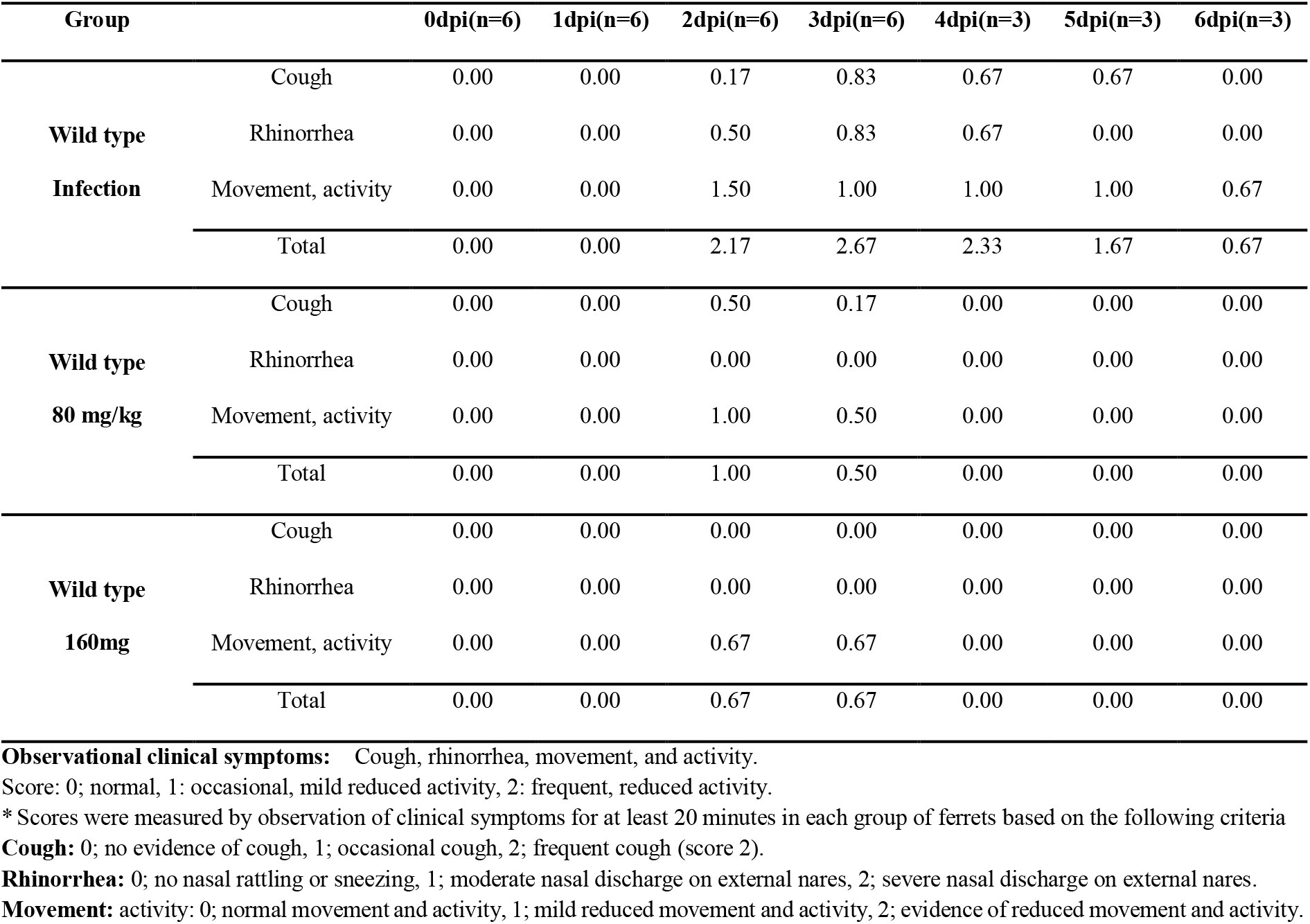
Clinical scores of ferrets infected with wild type SARS-CoV-2.

**Supplementary Figure 1.**
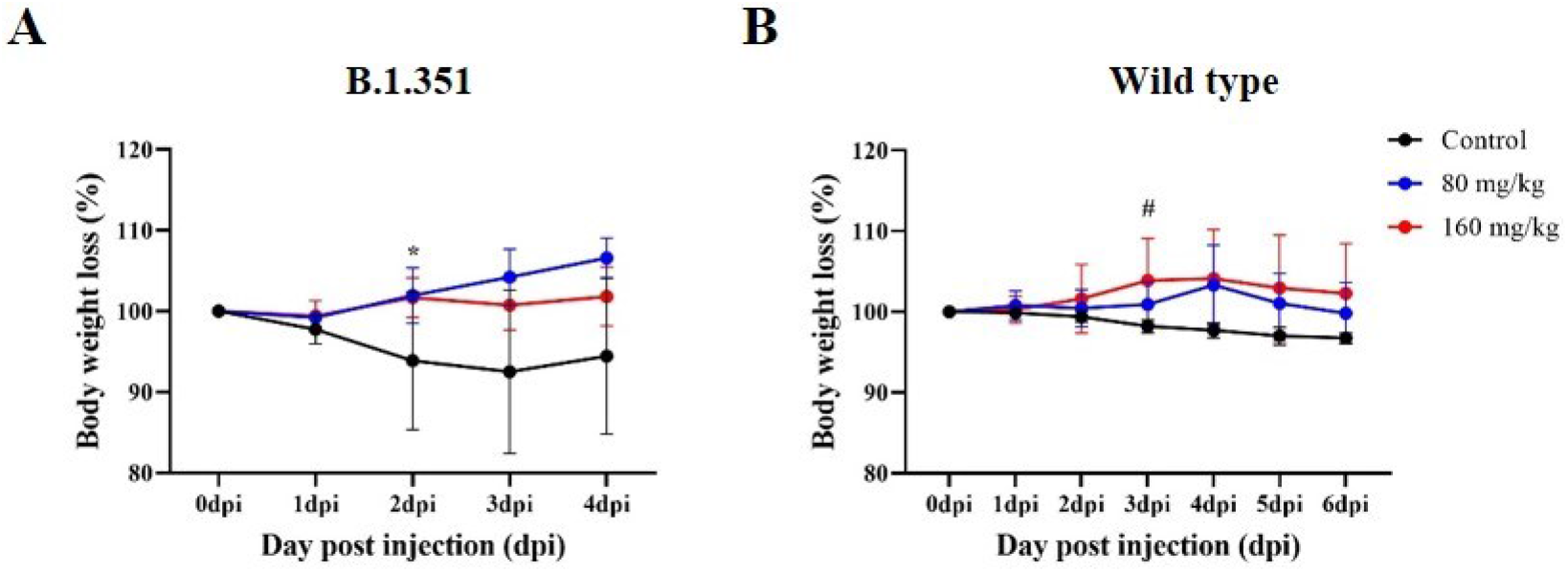
Ferret weight changes during *in vivo* efficacy evaluation of CT-P59. The body weights were daily measured for control group (vehicle) and CT-P59 treated groups (80 mg/kg and 160 mg/kg). The mean body weight changes were calculated based on individual baseline body weight from ferrets challenged with B.1.351 (A) and wild type virus (B). Asterisk or pound indicates statistical significance between the control and each group as determined by one-way ANOVA and post-hoc Dunnett’s test. (* indicates *P* < 0.05 in control vs 80 mg/kg and control vs. 160 mg/kg, # indicates P < 0.05 in control vs 160 mg/kg)

## Notes

### Competing Interest Statement

The authors have declared no competing interest.

